# In Diverse Conditions Intrinsic Chromatin Condensates Have Liquid-like Material Properties

**DOI:** 10.1101/2021.11.22.469620

**Authors:** Bryan A. Gibson, Claudia Blaukopf, Tracy Lou, Lynda K. Doolittle, Ilya Finkelstein, Geeta J. Narlikar, Daniel W. Gerlich, Michael K. Rosen

**Author notes:** Lead Contact. (M.K.R.).

## Abstract

Eukaryotic nuclear DNA is wrapped around histone proteins to form nucleosomes, which further assemble to package and regulate the genome. Understanding of the physical mechanisms that contribute to higher order chromatin organization is limited. Previously, we reported the intrinsic capacity of chromatin to undergo phase separation and form dynamic liquid-like condensates, which can be regulated by cellular factors. Recent work from Hansen, Hendzel, and colleagues suggested these intrinsic chromatin condensates are solid in all but a specific set of conditions. Here we show that intrinsic chromatin condensates are fluid in diverse solutions, without need for specific buffering components. Exploring experimental differences in sample preparation and imaging between these two studies, we suggest what may have led Hansen, Hendzel, and colleagues to mischaracterize the innate properties of chromatin condensates. We also describe how liquid-like in vitro behaviors can translate to the locally dynamic but globally constrained movement of chromatin in cells.

## INTRODUCTION

To maintain integrity during mitosis and fit into the nucleus, the eukaryotic genome must undergo substantial compaction (Olins and Olins, 2003). Across the cell cycle in eukaryotes each chromosome is spatially segregated from one another in dense structures, containing many loops (Cremer and Cremer, 2001). Individual loci of these chromosomes are constrained to move only within a locally defined region, controlled by interchromatin interactions, physical crosslinks induced by macromolecular complexes, and attachment of chromatin to static nuclear structures (Marshall et al., 1997; Mirny et al., 2019; Shaban and Seeber, 2020). A detailed account of the physical mechanisms that package the genome is critical, given the importance of spatial organization in regulating DNA templated processes such as transcription, DNA replication, and DNA repair (Dekker and Mirny, 2016; Finn and Misteli, 2019).

In a previous report we described the intrinsic capacity of chromatin to phase separate, producing liquid-like condensates with cell-like DNA density (Gibson et al., 2019). These intrinsic chromatin condensates can be regulated by cellular factors in kind with their functions in genome regulation (Gibson et al., 2019; Sanulli et al., 2019). We suggested interchromatin interaction through intrinsic condensation could represent a “ground state” for chromatin organization, molded or disrupted in cells by different regulatory factors (Boija et al., 2018; Eeftens et al., 2021; Gibson et al., 2019; Kagey et al., 2010; Lafontaine et al., 2021; Larson et al., 2017; Lee et al., 2021; Li et al., 2020; Plys et al., 2019; Sabari et al., 2018; Strom et al., 2017; Wang et al., 2019). A recent report from Hansen, Hendzel, and colleagues has called this work into question, suggesting that without specific buffering components intrinsic chromatin condensates are solid, reflecting the globally constrained organization of chromatin in cells (Strickfaden et al., 2020).

Here, we examine in detail the effect of solution conditions on the properties of intrinsic chromatin condensates. Across three different research groups using different chromatin sources and a variety of quantitative assays, we find that condensates composed of small chromatin fragments are fluid; a unique solution composition is not needed for their liquid-like properties. We also examine how experimental differences between these two reports may have given rise to the mischaracterization in Strickfaden et al. Cell 2020. Last, we make efforts to clarify how the liquid-like organization of condensates might translate to chromatin dynamics in cells.

## RESULTS

### BSA and DTT are Dispensable for the Liquid-like Properties of Intrinsic Chromatin Condensates

In Gibson et al Cell 2019, somewhat complex solutions were used to explore the nature of condensates formed from chromatin, most typically containing Tris buffer, acetate, potassium, magnesium, bovine serum albumin (BSA), dithiothreitol (DTT), ethylenediaminetetraacetic acid (EDTA), glycerol, and oxygen scavenging components (glucose oxidase, catalase, and glucose). The composition of this solution was an effort to mimic the cellular milieu (acetate, potassium, BSA, glycerol, and DTT) and reduce photodamage of condensates during fluorescence microscopy (oxygen scavenging components and DTT). In Strickfaden et al Cell 2020, the authors observed that in buffers lacking BSA and DTT condensates formed by a fluorescent dodecameric nucleosome array did not recover after photobleaching and did not fuse with one another. This led them to conclude that including BSA and DTT in buffers leads to artifactual liquid-like behavior of intrinsic chromatin condensates, and their omission reveals the true mesoscale material properties of condensates to be solid-like and constrained. In experimentation not reported in Gibson et al. Cell 2019 we had observed that neither BSA nor DTT were necessary for liquid-like chromatin condensates. To corroborate these observations, we set out to rigorously explore the effect of buffer conditions on chromatin condensate behavior.

We assembled dodecameric nucleosomal arrays by salt-mediated dialysis of reconstituted and unlabeled histone octamers and a DNA template with 12 repeats of Widom’s 601 nucleosome positioning element (Figure 1A). Using differential interference contrast microscopy, we observed in a minimal phase separation buffer composed of 25 mM Tris-acetate, 150 mM potassium acetate, and 1 mM magnesium acetate the formation of micron-sized spherical condensates that rounded upon fusion (Figure 1B) and maintained a consistent total volume following coalescence (Figure 1C).

**Figure 1.**
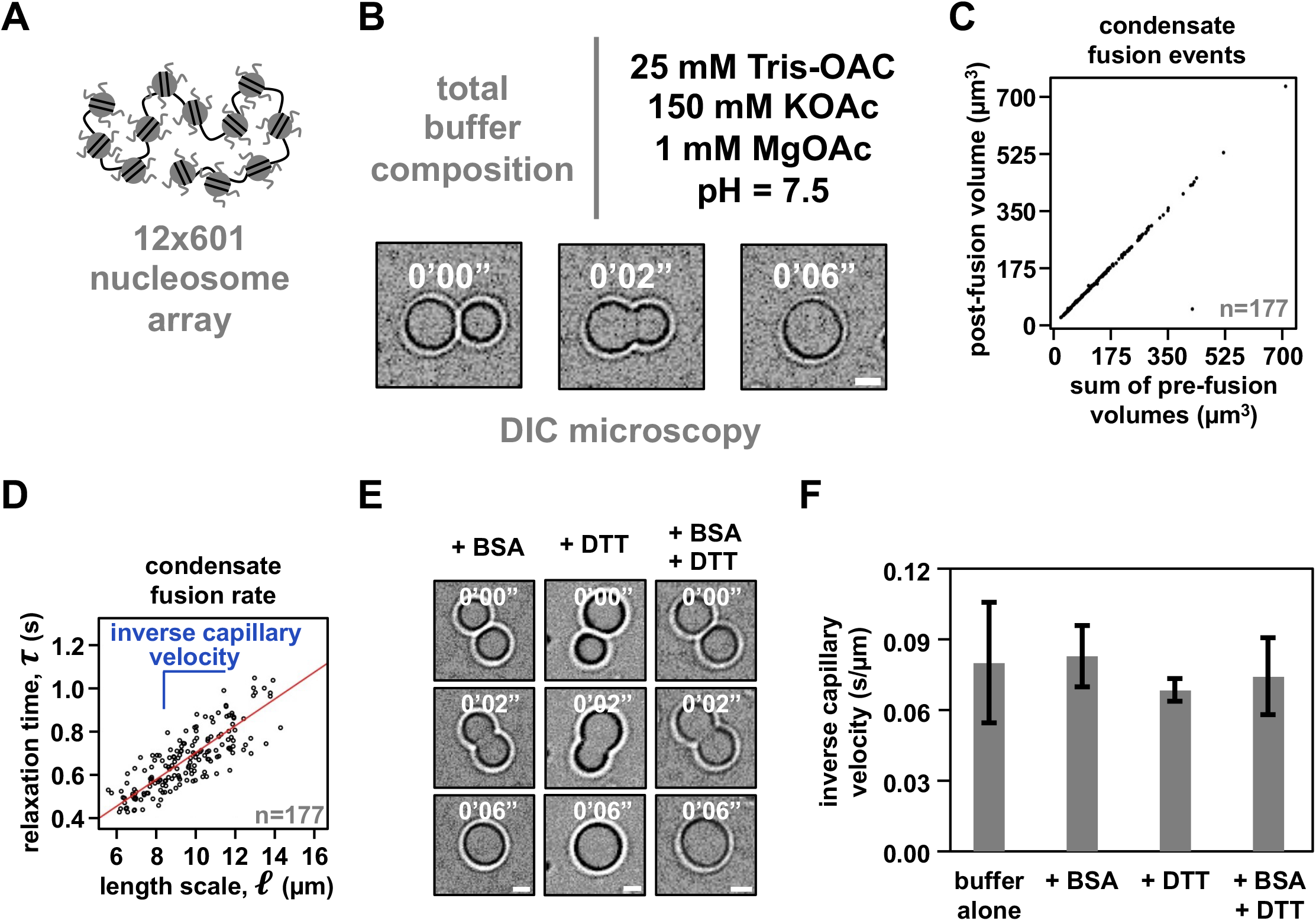
Intrinsic chromatin condensates are fluid without BSA and DTT. (A) Graphical depiction of the dodecameric nucleosomal arrays used for experimentation. (B) Differential interference contrast microscopy images of a fusion event between intrinsic chromatin condensates in the indicated buffer. (C) Dot plot representation of the inferred total volume of condensates before and after fusion. (D) Relaxation time versus length scale (sum of pre-fusion diameters) for 177 individual instances of condensate fusion in the buffer composition indicated in Figure 1B. Inverse capillary velocity, the characteristic ratio of surface tension, *γ*, and viscoscity, *η*, is derived from the linear fit (red line) of the plots’ slope. (E) Differential interference contrast microscopy images of intrinsic chromatin condensate fusion in the buffer indicated in Figure 1B supplemented with BSA (0.1 mg/mL, *left*), DTT (5 mM, *middle*), or BSA and DTT (0.1 mg/mL and 5 mM, respectively, *right*). (F) Bar chart of inverse capillary velocities (± standard deviation of 2 biological replicates) of intrinsic chromatin condensates in the buffer indicated in Figure 1B, buffer with BSA, or buffer with BSA and DTT. Scale bars, in white, are 4 µm.

Fusion of condensates followed by rounding to spherical shape is a hallmark of condensates with liquid-like material properties. The rate at which rounding occurs is a consequence of the relationship between the surface tension (*γ*) and viscosity (*η*) of condensates (Feric et al., 2016). Simple fluids coalesce according to the equation [inline], where ℓ is the diameter of condensates prior to fusion and *τ* is the characteristic relaxation time during coalescence. To determine *τ* for each instance of condensate fusion we measured the change in aspect ratio (*AR*) over time (*t*) during condensate fusion and found these values fit well to an exponential decay, *AR* = 1 + (*AR*_*init*_ − 1) · *e*^−t/τ^, where *AR*_*init*_ is the initial aspect ratio following the onset of fusion (Figure S1). Plotting *τ* versus ℓ from many fusion events (*N* = 177) showed clear linearity, with relaxation times on the order of seconds, indicating that intrinsic chromatin condensates in this minimal buffer are fluid (Figure 1D). The slope of this plot gives the inverse capillary velocity for these condensates in this solution, which is a quantitative measure of the distinctive ratio of surface tension (*γ*) and viscosity (*η*) of the material.

Intrinsic chromatin condensates formed in a solution containing BSA, DTT, or BSA and DTT also coalesced and became round (Figure 1E). The inverse capillary velocity was identical within error for condensates formed in minimal phase separation buffer alone, or buffer with BSA, DTT, or BSA and DTT (Figure 1F). These data do not support the conclusion in Strickfaden et al Cell 2020 that BSA and DTT are responsible for liquid-like material properties of intrinsic chromatin condensates.

### Intrinsic Chromatin Condensates are Liquid-like in a Variety of Solutions

We next explored how different anions and buffering systems affected the material properties of intrinsic chromatin condensates. Hansen, Hendzel, and colleagues state that “*liquid chromatin condensates were only observed under a single, highly specific set of conditions, requiring a combination of acetate anions, DTT, and BSA in addition to divalent cations. Under all other solution conditions tested, nucleosomal arrays were constrained and solid-like*”. We assayed the material properties of intrinsic chromatin condensates formed in solutions containing Tris buffer and sodium or potassium salts with chloride, acetate, or glutamate anions. Chloride is a typical anion used for biochemistry in buffered salt solutions. Previously, we used acetate to mimic small molecule anions in cells. And glutamate is the predominant anion found in cells (Park et al., 2016). We also used PIPES-KOH, a buffer/salt used in fluorescence-based assays that reconstitute cellular processes, including microtubule dynamics (Elie et al., 2015; Mitchison and Kirschner, 1984).

First, we determined the phase diagram for dodecameric nucleosomal arrays at 500 nM nucleosome concentration for each buffer (Figures 2A-2D). Condensates formed at similar concentrations of mono- and divalent salt in each buffering system, though glutamate anions required slightly higher concentrations of salt. In buffers containing chloride anion, condensate formation required at least 2 mM magnesium or the inclusion of glycerol (Figures S2A-S2F). While the source of this effect is not clear, it could arise from the propensity of glycerol to shield charged peptide side chains from salt (Mendes et al., 2003). Altogether, these data show that intrinsic chromatin condensation occurs robustly across many buffer compositions.

**Figure 2.**
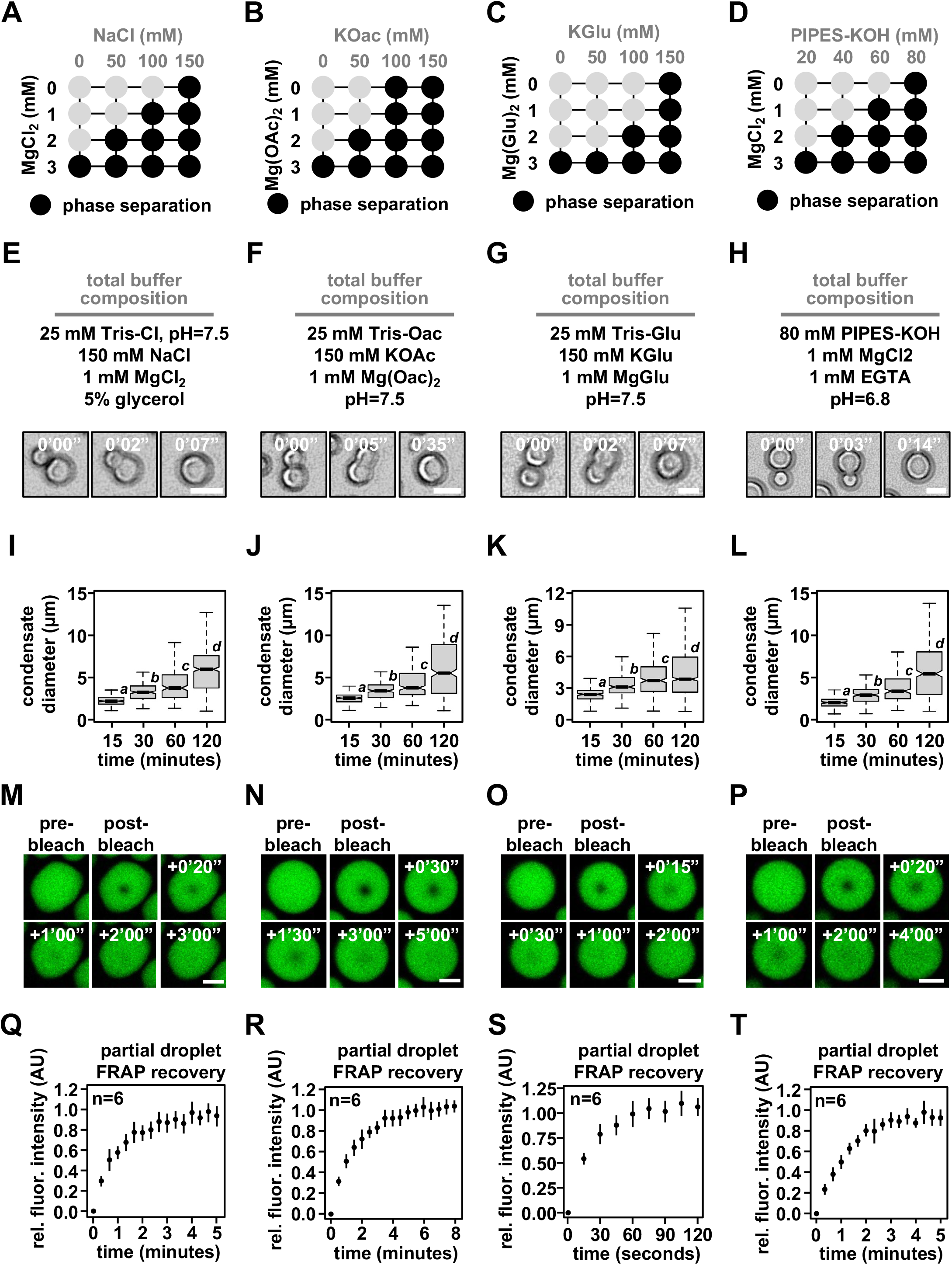
Intrinsic chromatin condensates are fluid in diverse buffers. Phase diagrams for intrinsic chromatin condensate formation in (A) Tris-chloride, (B) Tris-acetate, (C) Tris-glutamate, and (D) PIPES-KOH buffers. Dark Circles indicate the presence of condensates, and representative images are in Figure S2. Bright-field light microscopy images of intrinsic chromatin condensate fusion in (E) Tris-chloride, (F) Tris-acetate, (G) Tris-glutamate, and (H) PIPES-KOH buffers. Boxplots of intrinsic chromatin condensate diameters following induction of phase separation in (I) Tris-chloride, (J) Tris-acetate, (K), Tris-glutamate, or (L) PIPES-KOH based buffers. Bars marked with different letters are significantly different from one another (student’s t-test, p < 1×10^-7^). Fluorescence microscopy images of partial droplet FRAP of intrinsic chromatin condensates, in green, composed of nucleosomal arrays labeled with AlexaFluor 488 in (M) Tris-chloride, (N) Tris-acetate, (O), Tris-glutamate, or (P) PIPES-KOH based buffers. Quantification of partial droplet FRAP of intrinsic chromatin condensates in (Q) Tris-chloride, (R) Tris-acetate, (S), Tris-glutamate, or (T) PIPES-KOH based buffers. Fluorescence signal is normalized to post-bleach droplet intensity and error bars are standard deviation of 6 technical replicates. Scale bars, in white, are 4 µm.

For each buffering system, we chose a combination of mono- and divalent ions that resemble physiological concentrations in cells. In these solution conditions, both unlabeled (Figures 2E-2H) and AlexaFluor 488-labeled nucleosomal arrays (Figures S2G-S2J) rounded in seconds following fusion. Moreover, condensate size increased over the course of at least two hours (Figures 2I-2L), most likely through condensate fusion (Gibson et al., 2019). These data suggest that in different buffers intrinsic chromatin condensates are fluid. To probe the dynamics of molecules within these condensates, we photobleached a portion of condensates and measured the recovery of fluorescence (partial droplet FRAP) using condensates composed of AlexaFluor 488-labeled dodecameric nucleosomal arrays in each of the buffered salt solutions (Figures 2M-2T). These partial droplet photobleaching experiments were carried out using glass treatments that reduce condensate motion (see below) to aid in the quantitation of photobleach recovery. This preparation affects condensates in chloride buffers more strongly than others, resulting in adherence to the surface and non-spherical shapes. Still, in each buffer condition we observed rapid and full fluorescence recovery from photobleaching in minutes (Figures 2Q-2T). Notably, condensates in buffers with glutamate, the predominate anion in cells, recovered approximately three-fold more rapidly from photobleaching as compared to chloride, acetate, and PIPES-KOH buffered salt solutions (based on t1/2 of fluorescence recovery). These data refute the assertion in Strickfaden et al. Cell 2020 that liquid-like material properties are the result of specific conditions and show that in a variety of simple physiological salt solutions intrinsic chromatin condensates are fluid.

We note that in other solution conditions (Figures 2A-2D) the material properties of intrinsic chromatin condensates may differ. For example, in superphysiological concentrations of divalent salt alone, with no or minimal monovalent competitors, used historically to assess chromatin self-assembly (Hansen, 2002; Maeshima et al., 2016; Strickfaden et al., 2020), intrinsic chromatin condensates exhibit solid-like behaviors (Hansen et al., 2021). These condensates with reduced dynamics are likely the result of the extremely low ionic strength of the solution. How chromatin structures and dynamics in such non-physiologic conditions relate to biological systems remains to determined.

### Long Linker-length Chromatin with Complementary DNA Ends Forms Liquid Condensates

We next sought to understand differences in experimental conditions that might have led to mischaracterization of the properties of intrinsic chromatin condensates. A difference between Gibson et al. Cell 2019 and Strickfaden et al. Cell 2020 is the DNA template used to reconstitute chromatin. In Gibson et al. Cell 2019 chromatin was assembled on blunt-ended DNA with internucleosome linker DNA lengths from 15-45 base pairs. Hansen, Hendzel, and colleagues assembled chromatin on DNA with 4 bp palindromic single-stranded DNA overhangs and longer 60 base pair internucleosome linker DNA (Figure 3A). We prepared chromatin using the DNA template from Strickfaden et al. Cell 2020 and found that intrinsic chromatin condensates fused and rounded in seconds in minimal phase separation buffer lacking BSA or DTT, composed of 25 mM Tris-acetate, 150 mM potassium acetate, and 1 mM magnesium acetate (Figure 3B). This stands in contrast to observations in Strickfaden et al. Cell 2020 that chromatin condensates did not recover from whole droplet FRAP, and the resulting conclusion that the structures are solid, in these conditions (Figure 1J in (Strickfaden et al., 2020)). In partial droplet FRAP assays in the presence of either BSA or BSA and DTT these condensates each recovered in minutes within error of one another (Figures 3C-3E). These experiments demonstrate that altered material properties do not arise from differences in DNA template or an effect from BSA in the presence of DTT. This suggests that some facet of the experimentation as performed in Strickfaden et al. Cell 2020 led to solid-like behaviors from liquid-like condensates.

**Figure 3.**
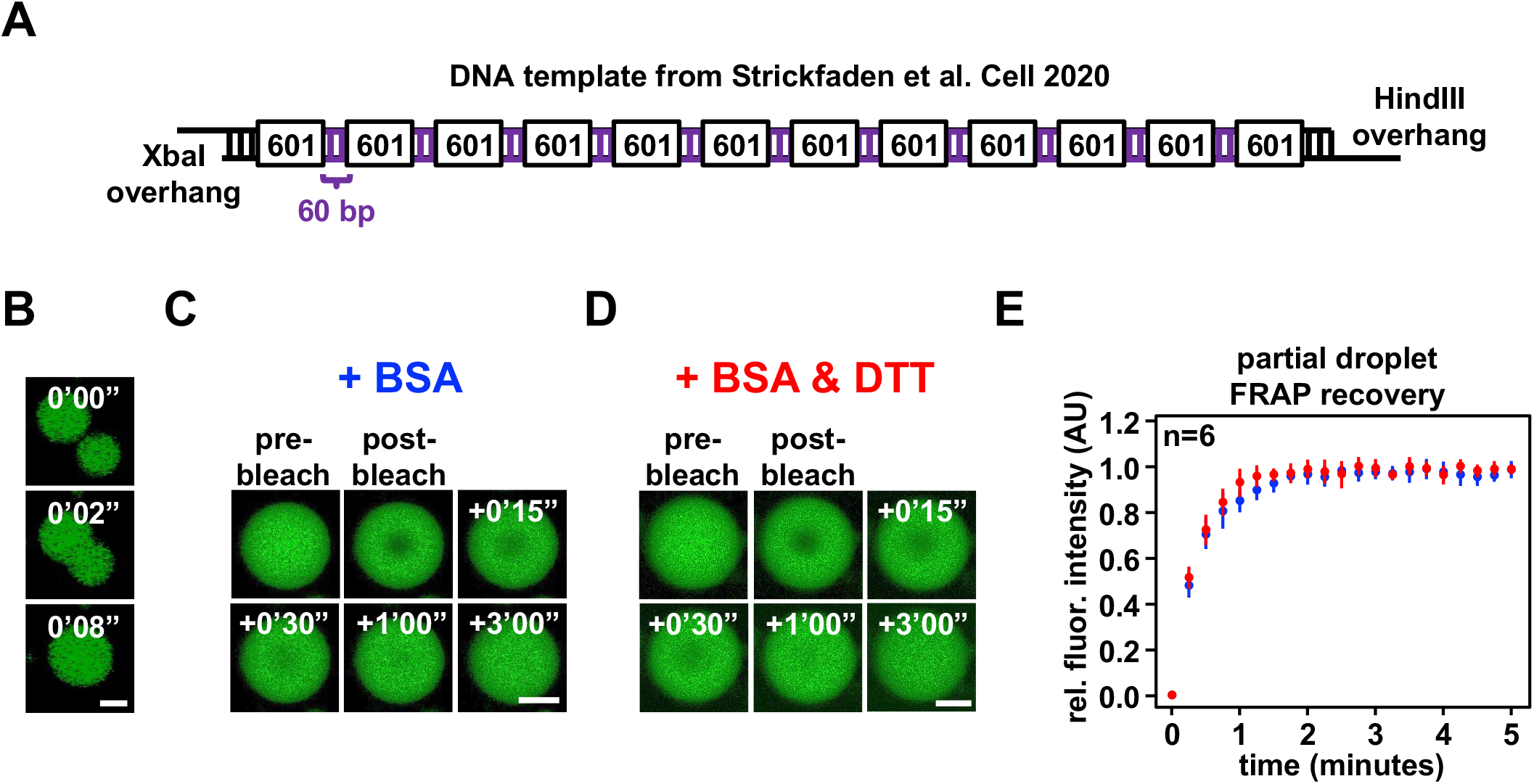
Chromatin reconstituted with the DNA template from Strickfaden et al Cell 2020 forms liquid-like condensates. (A) Graphical depiction of the 12×601 DNA template from Strickfaden et al Cell 2020. (B) Confocal Fluorescence Microscopy of intrinsic chromatin condensates composed of AlexaFluor 488-labeled nucleosomal arrays and the DNA template from Strickfaden et al. Cell 2020, in green, undergoing fusion. Confocal Fluorescence Microscopy of partial droplet FRAP of intrinsic chromatin condensates composed of AlexaFluor 488-labeled nucleosomal arrays and the DNA template from Strickfaden et al. Cell 2020, in green, formed in the presence of (C) 0.1 mg/mL BSA or (D) 0.1 mg/mL BSA and 5 mM DTT. (E) Quantification of partial droplet FRAP of intrinsic chromatin condensates with BSA or BSA and DTT, in blue and green, respectively. Fluorescence signal is relative to normalized to pre-bleach droplet intensity and error bars are standard deviation of 6 technical replicates. Scale bars, in white, are 4 µm.

### Sample Preparation Affects Condensate Movement and Exchange with Surroundings

In Strickfaden et al Cell 2020, chromatin condensates were deposited onto raw glass by centrifugation prior to fluorescence microscopy (Figure 4A). In Gibson et al. Cell 2019, we treated glass with PEG and BSA to prevent the adherence of macromolecules to the surface, as commonly done in biochemical imaging studies (Joo and Ha, 2012), and allowed condensates to settle onto the surface by gravity to minimize force-mediated perturbation (Figure 4B and Figure S1E of Gibson et al. Cell 2019). We investigated whether these differences affected the motion and physical properties of chromatin condensates.

**Figure 4.**
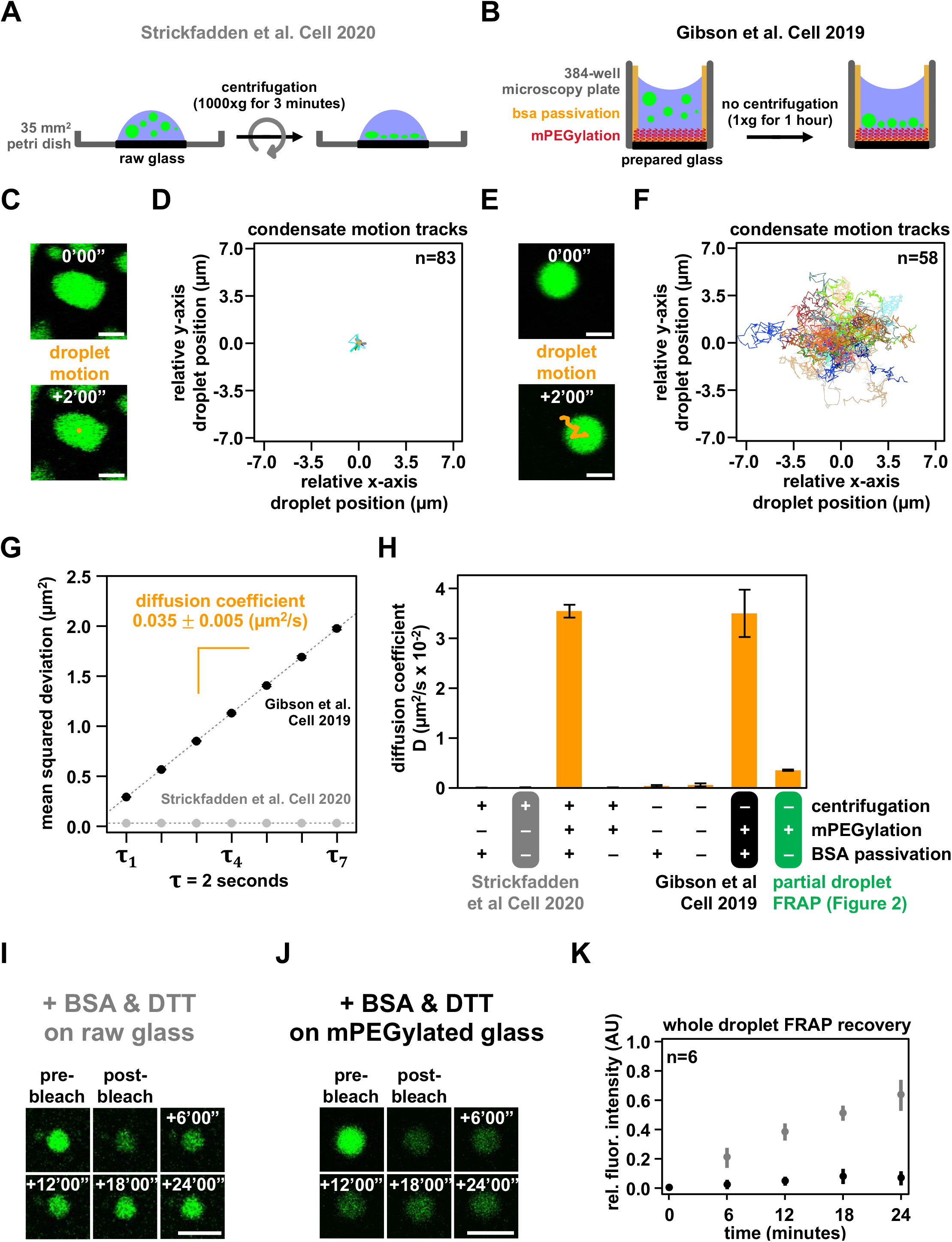
Condensate movement and dynamics is affected by microscopy glass preparation. Graphical depiction of techniques used to prepare intrinsic chromatin condensates for fluorescence microscopy imaging in (A) Strickfaden et al. Cell 2020 and (B) Gibson et al Cell 2019. (A) In Strickfaden et al. Cell 2020, intrinsic chromatin condensates were spun onto raw glass using a centrifuge. (B) In Gibson et al. Cell 2019, intrinsic chromatin condensates were added to 384-well microscopy plates and brought by gravity to rest on mPEGylated and BSA passivated glass. Movement of a single or many intrinsic chromatin condensates, following their preparation for fluorescence microscopy imaging as described in Strickfaden et al. Cell 2020 (C and D) and Gibson et al Cell 2019 (E and F). (C and E) The movement of an individual condensate across 2 minutes in 10 second intervals, are overlaid in orange on fluorescence microscopy images of AF488-labeled intrinsic chromatin condensates, in green. (D and F) The relative movement of many condensates determined across 2 minutes in 500 millisecond intervals. (G) Plot of mean squared displacement (± standard error) over lag time, *τ*, for intrinsic chromatin condensates prepared as described in Strickfaden et al. Cell 2020 (grey dots) and Gibson et al Cell 2019 (black dots). The diffusion coefficient, indicated in orange ± standard error, of intrinsic chromatin condensates can be calculated from the slope of the linear fit (dashed line) of the plotted data. (H) Bar chart of the diffusion coefficients of intrinsic chromatin condensates following their preparation for microscopy with and without centrifugation, mPEGylation of the microscopy glass, and BSA passivation of the microscopy well. Error bars are standard deviation of 4 technical replicates. Confocal fluorescence microscopy of whole droplet FRAP of intrinsic chromatin condensates composed of AlexaFluor 488-labeled nucleosomal arrays and the DNA template from Strickfaden et al. Cell 2020, in green, settled onto (I) raw glass or (J) mPEGylated glass. (K) Quantification of whole droplet FRAP recovery of intrinsic chromatin condensates on raw or mPEGylated glass, in grey and black, respectively. Fluorescence signal is relative to pre-bleach droplet intensity and error bars are standard deviation of 6 technical replicates. Scale bars, in white, are 4 μm.

Chromatin condensates deposited by centrifugation onto raw glass did not appreciably move during 2 minutes of observation by fluorescence microscopy, exhibited non-spherical morphology consistent with adhesion to the surface, and did not fuse because they could not contact each other (Figures 4C and 4D). In contrast, intrinsic chromatin condensates settled by gravity onto mPEGylated and BSA passivated glass moved many microns in distance, remained spherical and underwent fusion (Figures 4E-4F). We quantified movement in these two conditions by measuring the mean squared displacement by lag time and found that condensates settled onto prepared glass were mobile, with a diffusion coefficient of 0.035±0.005 µm^2^/s for condensates between 4-8 µm in diameter, while those deposited onto raw glass were not (Figures 4G, S3A-S3D, and S3K).

To understand what experimental parameter led to these differences, we quantified condensate movement with and without centrifugation, mPEGylation, and BSA passivation. Time-lapse imaging showed that diffusive condensate movement requires mPEGylation and BSA passivation, though some subdiffusive mobility is retained without passivation so long as glass is mPEGylated and condensates are not centrifuged onto the surface (Figures 4H and S3D-S3K). The microscopy sample preparation can thus impact condensate movement and fusion.

We considered whether BSA leaching from the passivated glass surface might lead to liquid-like condensate properties. Three pieces of data argue against this possibility. First, our photobleaching experiments, which show rapid recovery, are carried out in the absence of BSA passivation (Figures 2M-2T). Second, condensates move, albeit with restriction, in the absence of BSA passivation (Figure 3H). Third, condensates round following fusion with comparable kinetics in the presence or absence of BSA passivation (Figure S3L). Thus, the liquid-like behavior of intrinsic chromatin condensates is not a consequence of BSA passivation.

Given the strong effects of slide surfaces on condensate movement, we next asked how glass treatment might affect the physical properties of the condensates themselves. Using chromatin assembled with the DNA template from Hansen, Hendzel, and colleagues, we photobleached entire condensates to probe the extent of fluorescence recovery resulting from chromatin exchange between the condensed and dilute phases. This is distinct from partial droplet FRAP in Figure 3, which principally measures the movement of chromatin within a condensate. On raw glass, we observed appreciable recovery of fluorescence in the presence of BSA and DTT, consistent with results in Strickfaden et al. Cell 2020 (Figure 4I). In contrast, condensates settled onto mPEGylated glass did not substantially recover (Figures 4J and 4K) as we observed previously (Gibsone et al., Cell, 2019), a behavior we hypothesized is due to the very low concentration of chromatin in solution (note that differences in partial versus whole droplet FRAP recoveries were addressed in our previous study). These data show that microscopy preparations affect not just the movement of intrinsic chromatin condensates, but also their exchange with molecules in solution. While we do not understand the basis of this difference, condensates centrifuged to a strongly adherent glass surface will be flattened, perhaps appreciably so. In contrast, condensates settled onto a well passivated surface will remain spherical and shielded from the glass. The additional interactions between a flat condensate and glass may influence photobleaching recovery and might exhibit sensitivity to specific buffering components.

### BSA and DTT Mitigate Photo-crosslinking of Intrinsic Chromatin Condensates

Having analyzed how differences in sample preparation between these two studies alter condensate movement and FRAP recovery, we next examined potential differences in imaging. Laser excitation can produce radical oxygen species (ROS) that react with and crosslink neighboring molecules. Such light-induced crosslinking can cause artifactual hardening of biomolecular condensates (Ditlev et al., 2019). ROS production and photo-crosslinking of molecules are typically mitigated in biochemical imaging studies by including reducing agents in buffers, limiting fluorophore concentration, minimizing laser excitation, and scavenging soluble oxygen in solution (Dixit and Cyr, 2003; Joo et al., 2006; Zheng et al., 2014). The biochemical experiments of Strickfaden et al Cell 2020 did not include oxygen scavengers, raising the possibility that photo-crosslinking might have limited chromatin dynamics in their condensate imaging experiments. We therefore explored the effect of ROS mitigation on photo-crosslinking of intrinsic chromatin condensates.

We developed an assay to measure light-induced photo-crosslinking of intrinsic chromatin condensates. In this assay, condensates were formed in a buffer where free magnesium was required for their formation (Figure 2B, 2 mM Mg[OAc]2 and 50 mM KOAc). The concentration of monovalent salt in this buffer is insufficient to induce nucleosomal arrays to phase separate. Under these conditions, condensates can be dissolved by chelation of magnesium with EDTA (Figure 5A). We hypothesized that photo-crosslinking condensates would prevent their dissolution by EDTA.

**Figure 5.**
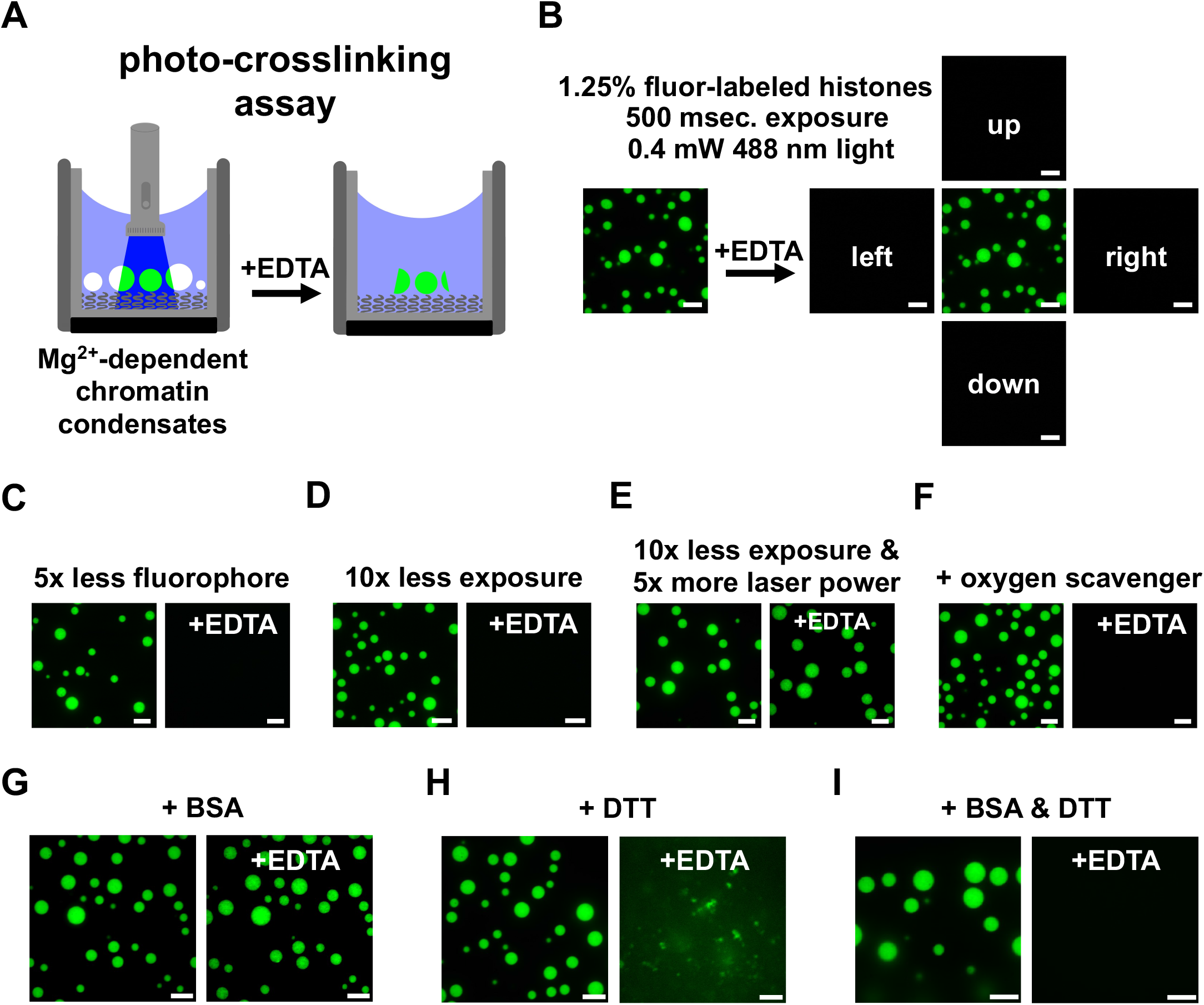
DTT and BSA mitigate photocrosslinking during fluorescence microscopy. (A) Diagram depicting an assay to detect photocrosslinking of intrinsic chromatin condensates. (*left*) Magnesium-dependent intrinsic chromatin condensates are exposed to fluorescent light prior to the addition of super-stoichiometric quantities of EDTA. (*right*) Photocrosslinked condensates fail to dissipate following chelation of magnesium. (B) Confocal fluorescence microscopy images of intrinsic chromatin condensates composed of nucleosomal arrays where 1 in 80 histone molecules are labeled with AlexaFluor 488. Images are following exposure to fluorescent light and both before (*left*) and after (*right*) the addition of EDTA. Confocal fluorescence microscopy images of intrinsic chromatin condensates imaged, as in Figure 5B, with (C) less fluorophore, (D) less exposure, (E) more laser power with less exposure, or (F) the inclusion of oxygen scavenging components. Confocal fluorescence microscopy images of intrinsic chromatin condensates formed in the presence of (G) BSA, (H) DTT, or (I) BSA and DTT and imaged as described in Figure 5B. Fluorescent microscopy images before and after the addition of EDTA were processed separately. Scale bars, in white, are 10 µm.

We formed intrinsic chromatin condensates in which 1 in 80 histone proteins are conjugated to a fluorophore (as in Strickfaden et al Cell 2020) in a magnesium-dependent phase separation buffer. Exposure of these condensates to 0.4 mW of fluorescent light (*λ* = 488 nm) prevented their dissolution by EDTA (Figure 5B). Condensates in adjacent fields, that had not been exposed to light, were dissolved 1 minute after the addition of EDTA. Light-induced solidification of condensates did not occur with five-fold less fluorophore or ten-fold less light (Figures 5C and 5D). Shorter exposure to light of higher intensity also led to condensate solidification, demonstrating that the totality and not duration of light exposure drives condensate solidification (Figure 5E). Addition of an oxygen scavenging system to the buffer prevents light-induced condensate solidification (Figure 5F), although its inclusion can alter condensate properties (Figure S4). Together, these data demonstrate that imaging intrinsic chromatin condensates can cause their solidification and suggests that this results from light-induced ROS production and photo-crosslinking. Furthermore, these data highlight how minimizing light exposure, fluorophore density, and including oxygen scavengers can prevent artifactual hardening of condensates.

We next sought to understand how the inclusion of BSA and/or DTT can influence photo-crosslinking of intrinsic chromatin condensates. Adding 100 ng/µL BSA, the concentration used in both Gibson et al Cell 2019 and Strickfaden et al Cell 2020, did not prevent condensate solidification (Figure 5G). In 5 mM DTT, light exposure and EDTA addition resulted in loss of spherical condensates but left aggregates in solution, suggesting partial but incomplete mitigation of photo-crosslinking (Figure 5H). Adding BSA and DTT together prevented condensate solidification, enabling their dissolution upon EDTA addition. While the mechanism by which BSA, or some component in commercially available BSA, can inhibit photo-crosslinking is unclear, these observations suggest that BSA and DTT can act in concert to reduce light-induced hardening of intrinsic chromatin condensates (Figure 4I). These data provide a plausible physical mechanism, imaging-induced solidification from ROS-mediated crosslinking, for the importance of BSA and DTT in condensate photobleach recovery reported by Hansen, Hendzel, and colleagues.

## DISCUSSION

### Intrinsic Chromatin Condensates are Liquid-like

Here, we present data acquired in three different laboratories, including some with independently prepared reagents (see Methods), demonstrating that intrinsic chromatin condensates are fluid under a wide range of solution conditions. Quantification of rounding after fusion and partial droplet FRAP recovery show that BSA and DTT impart no effect on condensate fluidity, even when using the DNA template of Hansen, Hendzel, and colleagues. From a series of experiments, we suggest that centrifugation of condensates onto raw glass and the omission of ROS-limiting components from buffers may have misled Hansen, Hendzel, and colleagues to characterize these condensates as solid-like. Our results have important implications on the behavior of chromatin and the use of phase-separated chromatin condensates to study nuclear processes.

### Bridging Fluid Condensates to Chromatin Dynamics in the Cell

A large body of data on the spatial organization and movement of loci in different cell types has demonstrated that on short length scales chromatin is highly dynamic. Super-resolution and single molecule fluorescence imaging studies have shown that nucleosomes compact into 30-50 nm clutches (Ricci et al., 2015), which further assemble into domains of ∼100-300 nm in radius (Ashwin et al., 2019; Itoh et al., 2021; Lakadamyali and Cosma, 2020; Nozaki et al., 2017b; Otterstrom et al., 2019). Analyses of their motion has shown that individual nucleosomes move within these domains on tens of milliseconds timescales (Gómez-García et al., 2021; Lerner et al., 2020), and the domains themselves move on hundreds of milliseconds to seconds timescales (Ashwin et al., 2019; Hajjoul et al., 2013; Itoh et al., 2021; Levi et al., 2005; Marshall et al., 1997; Nozaki et al., 2017b). In both regimes, movement is subdiffusive and/or confined (Ashwin et al., 2019; Gómez-García et al., 2021; Hajjoul et al., 2013; Itoh et al., 2021; Levi et al., 2005; Marshall et al., 1997; Nozaki et al., 2017a), in part due to constraints on a given chromatin segment imparted by adhesions to surrounding structures, which increase with length of the segment (i.e. number of adhesions) (Chubb et al., 2002; Hajjoul et al., 2013). Movement on these small scales is thought to primarily occur via passive thermal fluctuations rather than actively driven processes (Ashwin et al., 2019; Chubb et al., 2002; Hajjoul et al., 2013; Levi et al., 2005; Marshall et al., 1997; Nozaki et al., 2017b), although on longer timescales ATP-dependent jumps in position are also observed (Levi et al., 2005). Thus, short range/timescale movement reflect the dynamics of local internucleosome contacts that are subject to changes induced by histone acetylation and binding of linker histone H1 (Gómez-García et al., 2021; Otterstrom et al., 2019). These local dynamics are thought necessary for many genome functions, such as enhancer-promoter interactions (Chen et al., 2018), loop extrusion by SMC complexes (Davidson and Peters, 2021; Kim et al., 2019), and homologous pairing of sequences during meiosis and DNA repair (Hauer and Gasser, 2017; Kim et al., 2019). At greater scale (∼ 400 nm or larger) the genome does not appreciably move, constrained by the large size of chromosomes, crosslinking macromolecules (e.g., SMC complexes, adaptor proteins), and attachment of chromatin to nuclear structures (e.g., nuclear bodies, nuclear lamina) (Abney et al., 1997; Cremer et al., 1982; Gerlich et al., 2003; Marshall et al., 1997; Nozaki et al., 2017a). These constraints lead to the well-described reticence of chromatin in cells to recover from photobleaching (Abney et al., 1997; Hansen et al., 2017; Higashi et al., 2007; Kimura and Cook, 2001; Lever et al., 2000; Meshorer et al., 2006; Strickfaden et al., 2020). In condensates that form through interactions between small chromatin fragments alone, large-scale constraints are not present, allowing micrometer-scale movement and photobleach recovery. These long-range behaviors of intrinsic chromatin condensates in vitro very likely reflect the interactions that govern short length/time-scale chromatin dynamics in cells (Rippe, 2021).

To demonstrate how introducing one of these additional constraints, the very long length of chromosomes, would affect FRAP behavior we formed condensates composed of nucleosomal arrays with increasing length and found that photobleach recovery lessens at longer chromatin sizes (Figures 6A-6D). This length-dependent effect on condensate dynamics is general, as recently demonstrated with other biomolecular condensates in vitro (Keenen et al., 2021; Muzzopappa et al., 2021) or by fragmenting mitotic chromatin in cells (Schneider et al., 2021). This slowing of dynamics is driven by length-dependent steric occlusion of movement by neighboring molecules and increases in intermolecular contacts (Rubinstein and Colby, 2003). Nevertheless, even for very long polymers, short sections retain dynamics at short length scales while moving little at long length scales (Rubinstein and Colby, 2003). An intrinsic chromatin condensate composed of chromosome-length fragments would therefore be locally dynamic but exhibit little recovery from photobleach, like the dynamics of the genome observed in cells (Tortora et al., 2020).

**Figure 6.**
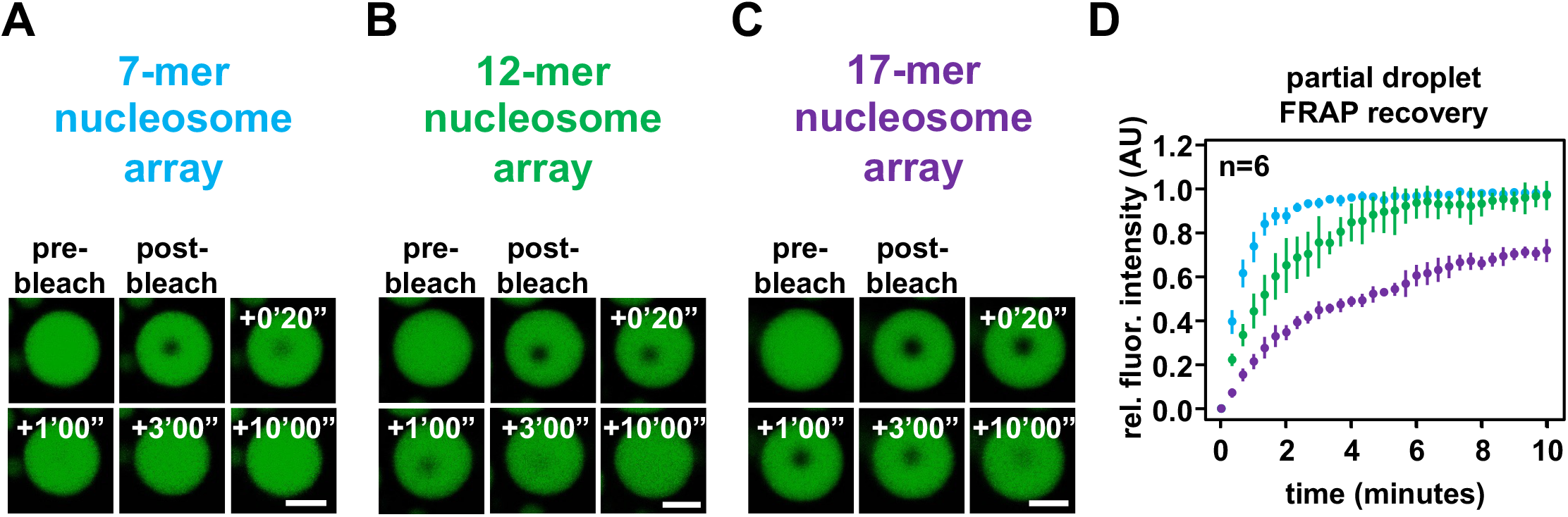
Length-dependent effects on chromatin condensate dynamics. Confocal fluorescence microscopy images of partial droplet FRAP of intrinsic chromatin condensates, in green, composed of AlexaFluor 488-labeled arrays that are (A) 7, (B) 12, or (C) 17 nucleosomes in length. (D) Quantification of partial droplet FRAP of intrinsic chromatin condensates composed of 7, 12, or 17 nucleosome-long arrays in blue, green, and purple, respectively. Fluorescence signal is relative to normalized pre-bleach droplet intensity and error bars are standard deviation of 6 technical replicates. Scale bars, in white, are 4 µm.

The FRAP recovery behaviors at different scales in Figure 6 underscores an important issue when studying condensates in vitro. Decades of study have demonstrated that the structure and function of discrete macromolecular complexes *in vitro* inform in a straightforward fashion on the structure and function of those factors *in vivo*. In contrast, the properties of condensates generated *in vitro* (e.g., size, structure, behavior) require care in their translation to cellular correlates. In this regard, we propose that factors that influence “mesoscale” genome dynamics in cells will not be readily observable when studying intrinsic chromatin condensates generated from kilobase scale DNA. Mesoscale genome dynamics, defined as the larger-scale motion that determines photobleach recovery of chromatin in cells, are likely governed by short-range chromatin interactions translated to genome-relevant scales in the context of complicating factors that crosslink and adhere chromatin to physical structures of the nucleus. The utility of the reconstituted system of phase-separated nucleosomal arrays is the ability to study how factors influence short-range chromatin dynamics using a macroscopic technique like FRAP.

## ACKNOWLEDGEMENTS

Research was supported by the Howard Hughes Medical Institute, a Paul G. Allen Frontiers Distinguished Investigator Award (to M.K.R.), grants from the NIH (R35 GM141736 to M.K.R., R35 GM127020 to G.J.N, F32GM129925 to B.A.G.), the Welch Foundation (I-1544 to M.K.R.), the European Research Council under the European Union’s Horizon 2020 research and innovation programme (grant agreement No 101019039, to D.W.G.), the Austrian Science Fund (SFB F34-06 and DK W1238 to D.W.G), the Wiener Wissenschafts-, Forschungs- und Technologiefonds (LS17-003 and LS19-001 to D.W.G), and the NSF (NSF-1921794 to G.J.N).

## AUTHOR CONTRIBUTIONS

M.K.R. and B.A.G. conceived of the study and designed the research program with D.W.G. and G.J.N. Using chromatin prepared in the Rosen laboratory, C.B. prepared material for microscopy and performed differential interference contrast microscopy appearing in Figure 1. In the Narlikar lab, T. L. assembled chromatin independently and performed bright-field microscopy appearing in Figure 2. L.K.D. and B.A.G. prepared biochemical materials used in the Rosen laboratory. B.A.G. performed all other in vitro biochemical experiments, fluorescence microscopy, and computational analyses. M.K.R., D.W.G., and G.J.N. secured funding and supervised the work. M.K.R. and B.A.G. wrote the manuscript with D.W.G. and G.J.N.

## SUPPLEMENTARY FIGURE LEGENDS

**Figure S1.**
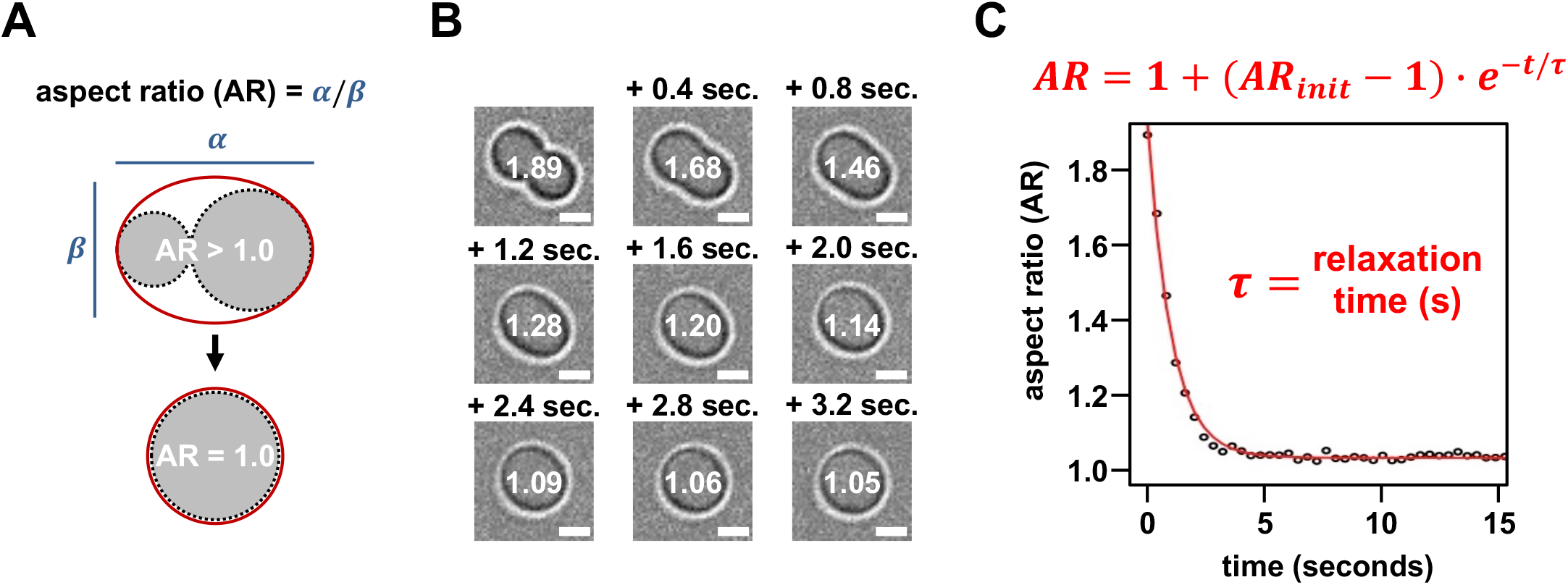
Quantitating condensate fusion rate. (A) Schematic depicting the calculation of aspect ratio (AR) during condensate fusion. (B) Time-lapse differential interference contrast microscopy of a rounding condensate following fusion along with the calculated aspect ratio determined for each acquisition. (C) Dot plot and exponential fit (red line) of a condensate fusion event from Figure S1B. Scale bars, in white, are 4 µm.

**Figure S2.**
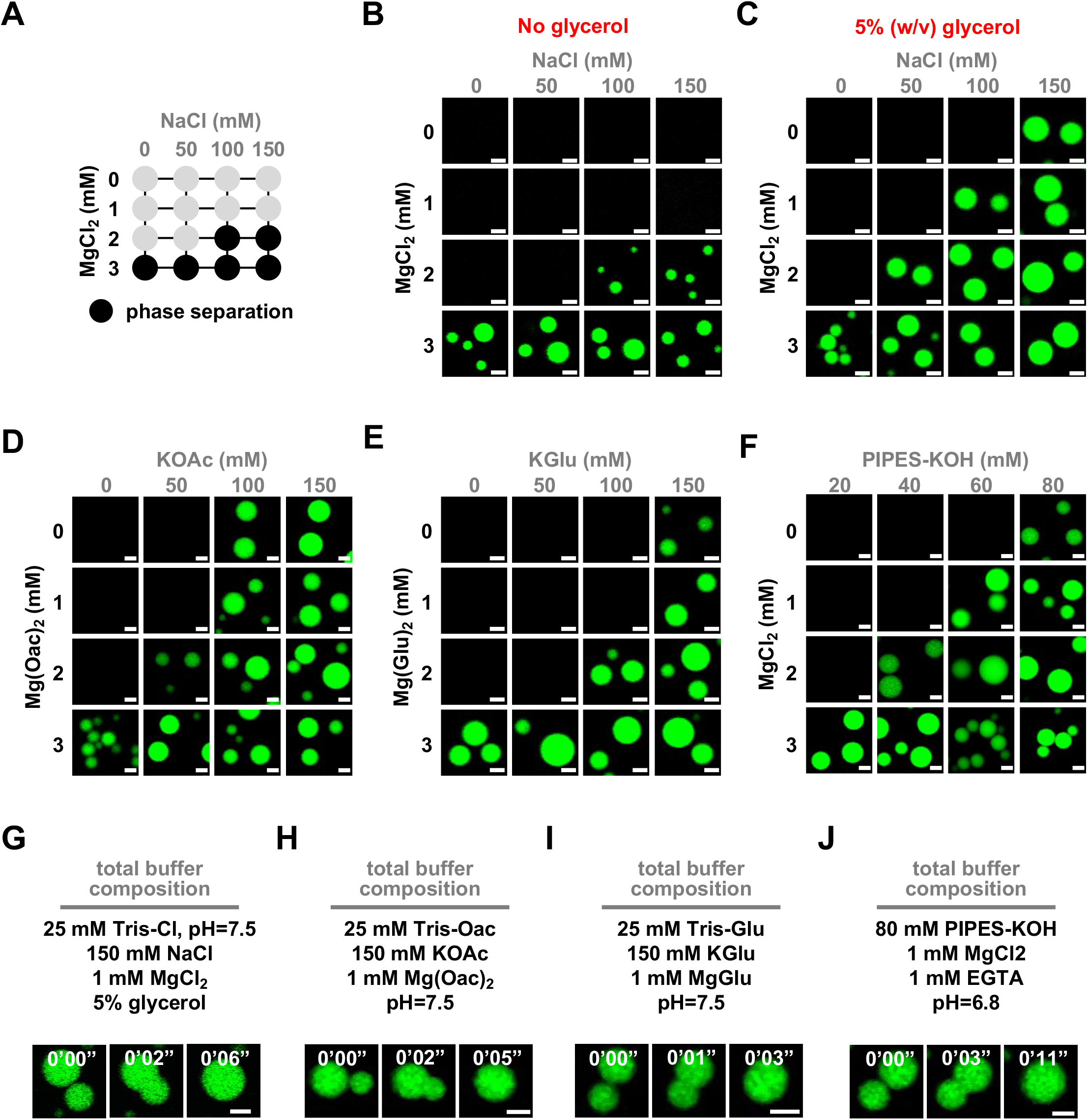
Phase separation of chromatin and fluor-labeled condensate fusion. (A) Phase diagram of chromatin in varying concentrations of NaCl and MgCl2 in Tris-chloride buffer without glycerol. Dark circles indicate the presence of condensates. Confocal fluorescence microscopy images of intrinsic chromatin condensates composed of AlexaFluor 488-labeled arrays, in green, in varying concentrations of salt in (B) Tris-chloride buffer without glycerol, (C) Tris-chloride buffer with glycerol, (D) Tris-acetate, (E) Tris-glutamate, and (F) PIPES-KOH. Time-lapse confocal fluorescence microscopy images of intrinsic chromatin condensates composed of AlexaFluor 488-labeled arrays, in green, undergoing condensate fusion in the indicated (G) Tris-chloride, (H) Tris-acetate, (I) Tris-glutamate, and (J) PIPES-KOH based buffered salt solutions. Scale bars, in white, are 4 µm.

**Figure S3.**
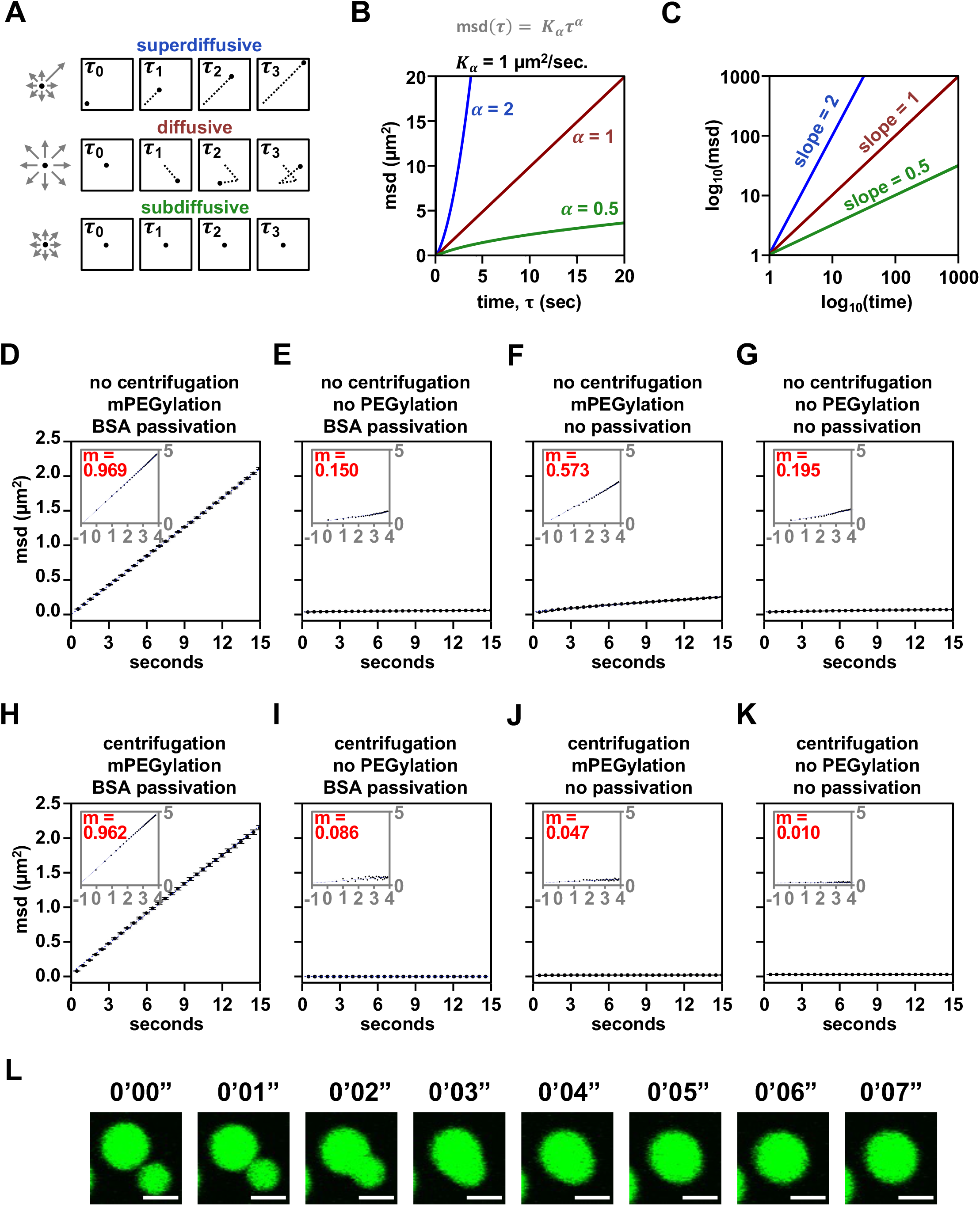
Motion of condensates with differing sample preparation and condensate fusion without BSA. (A) Graphical depiction of different kinds of particle motion. (B) Plot and (C) log-transformed plot of particle motion over time given different parameters for *K_α_* and *α*. (D-K) Plot of chromatin condensate motion over lag time with log transformation inset and slope of log-log plot in red. (L) Time-lapse confocal fluorescence microscopy images of intrinsic chromatin condensates composed of AlexaFluor 488-labeled arrays, in green, undergoing fusion without BSA passivation. Scale bars, in white, are 4 µm.

**Figure S4.**
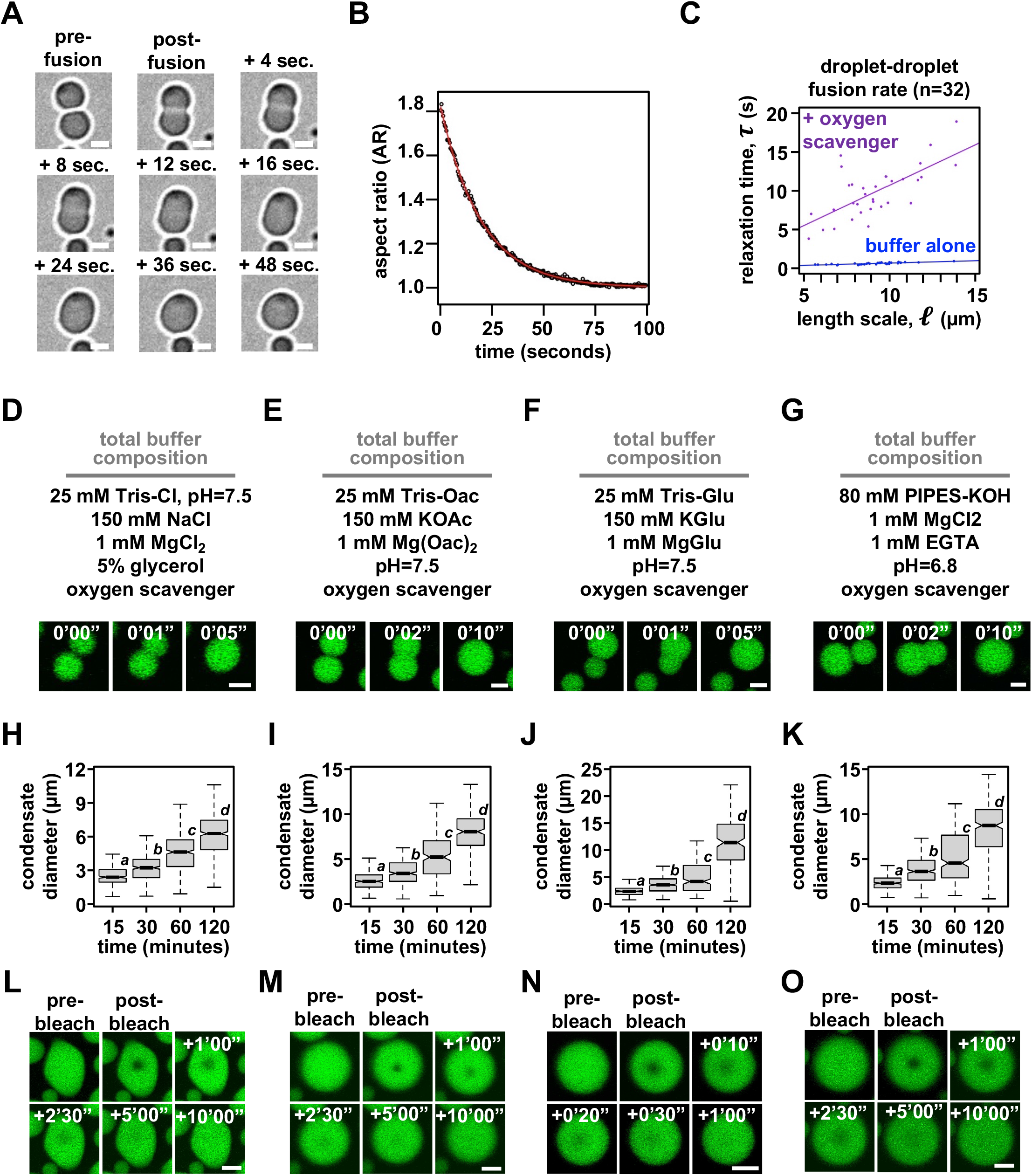
Condensate dynamics and properties in the presence of an oxygen scavenging system. (A) Time-lapse differential interference contrast microscopy of intrinsic chromatin condensates formed in the presence of oxygen scavenging components rounding after fusion. (B) Dot plot and exponential fit (red line) of a condensate fusion event from Figure S4A. (C) Relaxation time versus length scale (sum of pre-fusion diameters) for 32 individual instances of condensate fusion in the buffer composition indicated in Figure 1B, in blue, and that buffer containing oxygen scavenging components, in purple. Confocal fluorescence microscopy images of condensate fusion in (D) Tris-chloride, (E) Tris-acetate, (F) Tris-glutamate, and (G) PIPES-KOH buffers containing oxygen scavenging components. Boxplots of intrinsic chromatin condensate diameters following induction of phase separation in (H) Tris-chloride, (I) Tris-acetate, (J), Tris-glutamate, or (K) PIPES-KOH based buffers containing oxygen scavenging components. Bars marked with different letters are significantly different from one another (student’s t-test, p < 1×10^-7^). Fluorescence microscopy images of partial droplet FRAP of intrinsic chromatin condensates, in green, composed of nucleosomal arrays labeled with AlexaFluor 488 in (L) Tris-chloride, (M) Tris-acetate, (N), Tris-glutamate, or (O) PIPES-KOH based buffers containing oxygen scavenging components. Scale bars, in white, are 4 µm.

## STAR METHODS

### Lead Contact and Materials Availability

Further information and requests for resources and reagents should be directed to and will be fulfilled by the Lead Contact, Michael K. Rosen.

### Experimental Model and Subject Details

#### Bacterial Strains

DH5α (Invitrogen) and MACH1 (Invitrogen) *E. coli* strains were used for passage during cloning of plasmid DNA. Large-scale preparations of plasmid DNA for isolation of nucleosome assembly sequences were passed through and grown to scale in the ER2925 (*dam^-^/dcm^-^*) *E. coli* strain (NEB). Histone proteins were expressed in Rosetta2(pLysS) E. coli cells (Novagen) in the Rosen lab and in BL21(DE3) pLysS cells (Agilent) in the Narlikar lab.

### Method Details

#### Molecular Biology and Cloning

##### Construction of 12×601 dsDNA Array-Producing Bacterial Vectors

###### 193bp Repeat Length TetO-containing 12×601

The p12×601 insert from Gibson et al Cell 2019 was subcloned into the WM530 plasmid (a generous gift from Tom Muir) to create pWM+12×601 plasmid. This plasmid contains a 12×601 array with Tet Operator (TetO) inserted between 601 sequences 6 and 7 with 46 bp DNA lengths between nucleosome positioning sequences. Digesting this plasmid with EcoRV fragments the DNA backbone into < 500 bp sizes (e.g., carrier DNA) and liberation of the 12×601 array.

###### 7×601 and 17×601 Nucleosomal Arrays

pUC19 with a 7×601 insert containing 25 base pair DNA segments between nucleosome positioning sequences was subcloned from pUC19+6×601_25bplinker (Gibson et al., 2019) and pUC19+3×601_25bplinker plasmids (Gibson et al., 2019) to create pUC19+7×601_25bplinker. pUC19 with a 17×601 insert containing 25 base pair DNA segments between nucleosome positioning sequences was subcloned from pUC19+7×601_25bplinker and pUC19+12×601_25bplinker (Gibson et al., 2019) plasmids to create pUC19+17×601_25bplinker. Of note: 2 601 repeats are lost during isoschizomer digestion and ligation when subcloning these 601-repeat containing constructs. 601 array DNA within pUC19+7×601_25bplinker and pUC19+17×601_25bplinker were subcloned into WM530 to create pWM+7×601 and pWM+17×601. Digesting this plasmid with EcoRV fragments the DNA backbone into < 500 bp sizes (e.g., carrier DNA) and liberation of the 12×601 array.

#### Expression and Purification of Recombinant Proteins in the Rosen Lab

##### Purification of H. sapiens Histone Proteins Expressed in E. coli

###### Expression

Histones were expressed exactly as previously described (Gibson et al., 2019). Briefly, pET-based plasmids encoding human histone proteins (H3C111A, H4, H2A.1, H2B, or H2BT116C) were transformed into Rosetta2(pLysS) E. coli cells (Novagen) and gorwn to a density (OD600) of 0.4 at 37°C. Recombinant protein expression was induced by addition of IPTG to 1 mM for 3 hours at 37°C. The cells were collected by centrifugation, resuspended in Histone Lysis Buffer (50 mM Tris•HCl, pH 8, 150 mM NaCl, 5 mM ß-mercaptoethanol, 1 mM Benzamidine, 100 µM Leupeptin, 100 µM Antipain, 1 µM Pepstatin), flash frozen in liquid N2, and stored at −80°C.

###### Purification

Histones were purified essentially as previously described (Gibson et al., 2019). *E. coli* expressing histone proteins were passed through an Avestin Emulsiflex-C5 high pressure homogenizer at ∼10,000 PSI. Inclusion bodies were washed twice by centrifugation and resuspension with Inclusion Body Wash Buffer (50 mM Tris•HCl, pH 7.5, 100 mM NaCl, 1% Triton X-100, 1 mM EDTA, 1 mM Benzamidine, 5 mM ß-mercaptoethanol) and twice more omitting Triton X-100. Inclusion Bodies were soaked with DMSO, minced with a spatula, extracted with Histone Unfolding Buffer (20 mM Tris•HCl, pH 7.5, 7M Guanidinium-HCl, 10 mM DTT). Extracted unfolded proteins centrifuged, filtered through a 0.45 µm membrane (GE Healthcare) and run in Histone Unfolding Buffer over a HiLoad 26/60 Superdex 200 pg size exclusion column. Fractions containing histone proteins were dialyzed three times against 5 mM ß-mercaptoethanol. Any precipitate was pelleted by centrifugation.

Soluble histone protein was reduced in volume in a centrifugal concentrator before dilution with > 20 volumes of SAU200 (20 mM NaOAc, pH 5.2, 7 M Urea, 200 mM NaCl, 1 mM EDTA, 5 mM ß-mercaptoethanol). Histone proteins in SAU200 were filtered and purified by cation exchange chromatography (Source 15S) using SAU200 (20 mM NaOAc, pH 5.2, 7 M Urea, 600 mM NaCl, 1 mM EDTA, 5 mM ß-mercaptoethanol) as an eluate. Fractions containing histone proteins were dialyzed three times against 5 mM ß-mercaptoethanol, reduced in volume in a centrifugal concentrator, quantified by measuring solution absorbance at 280 nm and using the calculated molar extinction coefficients (https://web.expasy.org/protparam/) for histones H3C111A, H4, and H2A of 4470/M•cm, 5960/M•cm, and 4470/M•cm, respectively. Purified histone proteins were aliquoted in single use quantities, flash frozen with liquid N2, and stored at −80 °C.

##### Labeling Histone H2BT116C with AlexaFluor 488

Histone H2B mutant H2BT116C was labeled exactly as previously described (Gibson et al., 2019). Briefly, cysteines reduced by adding TCEP to 1 mM final concentration at room temperature for 1 hour. Following salt exchange of Histone H2BT116C into Phosphate Buffered Saline (8 mM Na2HPO4, 2 mM KH2PO4, pH 7.4, 137 mM NaCl, 2.7 mM KCl) using HiTrap Desalting Columns (GE Healthcare) 1.5 molar excess Alexa Fluor 488 (AF488)-C5-maleimide was added. Following a 4-hour incubation in the dark, DTT was added to 10 mM final concentration to quench the reaction. Free fluorophore was removed by passing fluor-conjugated histone H2B a desalting column and anion exchange chromatography (Source 15Q) in Histone CleanUp Buffer (20 mM Tris•HCl, pH 7.5, 150 mM NaCl, 1 mM DTT). Fractions containing AF488-labeled histone H2B proteins were dialyzed three times against 5 mM ß-mercaptoethanol and reduced in volume using centrifugal concentrators. Protein concentration and percent labeling were quantified by measuring absorbance at 280 and 495 nm and the calculated molar extinction coefficients (https://web.expasy.org/protparam/ or Thermo Scientific) for histone H2BT116C and AlexaFluor488 of 4470/M•cm and 73000/M•cm, respectively. 100% labeling was confirmed and AlexaFluor 488-labeled histone H2B protein was aliquoted for single use, flash frozen with liquid N2, and stored at −80°C.

#### Expression and Purification of Recombinant Proteins in the Narlikar Lab

*Xenopus laevis* histones were expressed in BL21(DE3) pLysS cells (Agilent) and purified from *E.coli* as previously described (Luger et al., 1999).

#### Reconstitution of Histone Octamers in the Rosen Lab

Histone Octamers were reconstituted essentially as previously described (Gibson et al., 2019). Briefly, aliquots of histone proteins were mixed in Histone Unfolding Buffer at a final concentration of 16.7:16.7:20:20 nmol per mL for H4:H3:H2B:H2A. The histone mix was dialyzed three times against Refolding Buffer (10 mM Tris•HCl, pH 7.5, 2 M NaCl, 1 mM EDTA, 5 mM ß-mercaptoethanol). The dialysate was filtered by passage through a 0.45 µm filter and reduced in volume using a centrifugal concentrator. Refolded histone octamer was isolated by size exclusion chromatography on a HiLoad SD200 26/60 pg column. Peak Fractions were analyzed by SDS-PAGE analysis for equal stoichiometry of core histone proteins, pooled, and reduced in volume with a centrifugal concentrator. The absorbance of reconstituted histone octamers was measured at 280 nm, (and 495 nm for fluorophore-labeled octamers) and concentration of protein was calculated using the molar extinction coefficients (https://web.expasy.org/protparam/ or Thermo Scientific) for histone octamer and AF488 of 44700/M•cm and 73000/M•cm, respectively. 100% labeled histone octamers were confirmed by the presence of 2:1 stoichiometric excess of fluor to histone octamer. Purified histone octamers were aliquoted, flash frozen with liquid N2, and stored at −80 °C. Note: The final concentration of histone octamers used for chromatin assembly was adjusted by their capacity to assemble single 601 sequences into mononucleosomes.

#### Reconstitution of Histone Octamers in the Narlikar Lab

Histones were refolded in high salt buffer to form octamer and purified by size-exclusion chromatography as previously described (Luger et al., 1999).

#### Preparation of DNA templates for Chromatin Assembly in the Rosen Lab

Plasmids containing 601 repeat DNA were prepared largely as previously described (Gibson et al., 2019). Briefly, plasmids containing 601 repeat DNA were transformed into dam^-^/dcm^-^ *E. coli* strain ER2925 and plated onto LB agar plates supplemented with 100 ng/µL Ampicillin for growth overnight. Following small scale growth to turbidity from a single colony, 4.5 liters of LB with 100 ng/µL of Ampicillin were inoculated with bacteria for overnight growth at 37 °C. Bacteria was harvested by centrifugation and plasmid DNA was purified using a Qiagen Plasmid Giga Kit. 601 array DNA was liberated from the plasmid backbone and backbone DNA digested to small fragments (e.g., carrier DNA) using restriction endonuclease digestion. pWM+12×601, pWM+12×601_25bplinker, and pWM+17×601_25bplinker were digested with the EcoRV-HF restriction endonuclease. pWM+7×601_25bplinker was digested with EcoRV-HF, MSPA1I, TaqI-V2, and AvaII restriction endonucleases. p12×207, used in Strickfaden et al. Cell 2020 (A generous gift from Jeffrey Hansen), was digested with XbaI, HindIII, MspA1I, AatII, BsaWI, DraI, and BglI. Following backbone digestion and 601 array DNA liberation, DNA was purified by Phenol:Chloroform:Isoamyl Alcohol (25:24:1) Extraction and Ethanol Precipitation. DNA was resuspended for storage in 1xTE (10 mM Tris•HCl, pH 7.5, 1 mM EDTA) and quantified using a Nanodrop (ThermoFisher Scientific).

#### Preparation of DNA templates for Chromatin Assembly in the Narlikar Lab

The array DNA template was isolated by restriction enzyme digest of plasmid containing 12×601s (Gibson et al., 2019). Array DNA was purified using the Gigaprep Kit (Qiagen) and by size-exclusion. This was followed by ethanol precipitation and resuspension in 1X TE.

#### Preparation of Nucleosomal Arrays in the Rosen Lab

##### Setup of Nucleosomal Assemblies

Nucleosomes were setup for assembly largely as previously described (Gibson et al., 2019). Briefly, DNA and histone octamers were thawed on wet ice. An equal volume of 4M Assembly Buffer (10 mM Tris•HCl, pH 7.5, 1 mM EDTA, 4 M NaCl, 2 mM DTT) was added to DNA (in 1xTE). Histone octamers were added in slight stoichiometric excess relative to 601 nucleosome positioning sequences in the template (1.3:1), to ensure full assembly in the presence of carrier DNA which prevents overassembly. Final concentrations of octamer/601 was 5 µM. Histone octamers and 601-containing DNA templates were moved into dialysis chambers equilibrated in High Salt Assembly Buffer (10 mM Tris•HCl, pH 7.5, 1 mM EDTA, 2 M KCl, 1 mM DTT).

##### Salt Dialysis-mediated Assembly of Nucleosomes

Nucleosomes were assembled as previously described (Gibson et al., 2019). The salt concentration in dialysis chambers containing histones and DNA were lowered over three days by continuous dilution using Low Salt Assembly Buffer (10 mM Tris•HCl, pH 7.5, 1 mM EDTA, 200 mM KCl, 1 mM DTT) at 4°C. After three days dialysis chambers were dialyzed against 1xTE with 1 mM DTT for at least for hours at 4°C.

##### Sucrose Gradient-mediated Purification of Nucleosomal Arrays

Following salt-mediated dialysis, assembled nucleosomal arrays were applied to linear 15-40% (12- or 17-nucleosome arrays) or 5-25% (7-nucleosome arrays) sucrose gradients in 1xTE with 1 mM DTT. Sucrose gradient fractions containing assembled nucleosomal arrays were dialyzed into 1xTE with 1 mM DTT and concentrated in using centrifugal concentrators.

##### Quantitation of Nucleosome Concentration

To quantitate final chromatin concentrations, 10 µL of nucleosomal arrays were added to 90 µL of SDS/PK Buffer (45 mM Tris•HCl, pH 7.5, 9 mM EDTA, 1% SDS) and incubated for 30 minutes at room temperature. DNA was purified using a Qiagen PCR purification Kit and quantified using a Nanodrop spectrophotometer.

##### Quality Assurance of Nucleosomal Assembly

The quality of nucleosome assemblies was assessed for by digesting linker DNA between 601 sequences in template DNA and chromatin using a restriction endonuclease for 1 hour at room temperature and running the digests on a native PAGE gel in 0.5X TAE. Only chromatin without unassembled 601 sequences, a clear shifted mononucleosome band, and at most a trace hexasome population were used for experimentation.

#### Preparation of Nucleosomal Arrays in the Narlikar Lab

12-mer nucleosome arrays were generated from a salt gradient dialysis to assemble histone octamer onto the 12×601 DNA as previously described (Sanulli et al., 2019). After assembly, arrays were dialyzed into TCS0.1 (20mM HEPES pH7.5, 0.1mM EDTA, 2mM 2-Mercaptoethanol).

#### Preparation of 384-well Microscopy Plates in the Rosen Lab

##### mPEGylation of Silica

384-well microscopy plates (Brooks Life Science Systems Matriplate) were washed with 5% Hellmanex at 37°C for 4 hours and then rinsed copiously with ≥ 18 MΩ H2O. Silica was etched with 1 M NaOH for 1 hour at room temperature and then rinsed copiously with ≥ 18 MΩ H2O. Depolymerized Silica was covalently bonded overnight (≥ 18 hours) at room temperature to 20 mg/mL 5K mPEG-silane (PEGWorks) suspended in 95% Ethanol. Plate was washed many times with 95% Ethanol, rinsed with copious amounts of ≥ 18 MΩ H2O, and completely dried in a chemical hood over 3-4 hours. PEGylated microscopy plate was sealed until individual wells’ use with an adhesive PCR plate foil (Thermo).

##### Passivation of Well with Bovine Serum Albumin

Following PEGylation, foil was cut above individual wells prior to their use and both plastic and PEGylated glass were passivated by incubation with freshly prepared 100 mg/mL BSA for 30 minutes to 4 hours. Wells were rinsed three times with ≥ 18 MΩ H2O to remove BSA and microscopy samples (15-40 µL in volume) were immediately added to the empty well. Desiccation of microscopy samples was limited following their addition to the plate by sealing with transparent scotch tape.

#### Preparation of 384-well Microscopy Plates in the Narlikar Lab

##### mPEGylation of Silica

Wells in a microscopy plate were washed twice with ddH2O before adding 2% Hellmanex. After a 1-hour incubation at room temperature the Hellmanex solution was removed, and wells were washed three times with ddH2O. 0.5 M NaOH was added to the wells to etch the glass for 30 minutes at room temperature. The NaOH solution was removed and wells were washed three times with ddH2O before adding 20 mg/mL mPEG-silane dissolved in 95% Ethanol. Wells were covered with foil tape and left to sit overnight in the dark. Nearly all of the mPEG-silane was removed before 10 serial washes with 95% EtOH followed by three washes with ddH2O. Through each wash trace amounts of liquid were retained in the well to prevent dessicating the PEG layer conjugated to glass.

##### Passivation of Well with Bovine Serum Albumin

Following PEGylation, 100 mg/mL BSA was added to passivate the well for 2 hours at room temperature. Wells were rinsed three times with ddH2O, finally leaving water in the well to await transfer of phase-separated chromatin.

#### Preparation of 384-well Microscopy Plates in the Gerlich Lab

##### mPEGylation of Silica

384-well µClear^®^ microscopy plates (Greiner Bio-One, 781906) were washed with 5% Hellmanex in ≥ 18 MΩ H2O at 65°C for 4 hours in a tabletop Incu-Line oven (VWR) then rinsed 10 times with ≥ 18 MΩ H2O. Silica was etched with 1 M KOH for 1 hour at room temperature and then rinsed 10 times with ≥ 18 MΩ H2O. The etched multi-well plate was treated with 5K-mPEG-silane (Creative PEGWorks, PLS-2011) suspended in 95% Ethanol (VWR) for 18 hours at room temperature. The plate was washed once with 95% Ethanol, then 10 times with ≥ 18 MΩ H2O, before completely drying in clean chemical hood overnight. Until their use, wells were sealed using an adhesive PCR foil (Thermo Fisher Scientific) and kept in a dry dark space.

##### Passivation of Well with Bovine Serum Albumin

Following PEGylation, foil was cut above individual wells prior to their use and both plastic and PEGylated glass were passivated by incubation with freshly prepared 100 mg/mL BSA for 30 minutes to 4 hours. Wells were rinsed three times with ≥ 18 MΩ H2O to remove BSA and microscopy samples were immediately added to the empty well. Desiccation of microscopy samples was limited following their addition to the plate by sealing with transparent scotch tape.

#### Imaging Condensates in the Rosen Lab

##### Phase Separation of Nucleosomal Arrays

Nucleosomal arrays with a 25 bp internucleosome linker DNA length and 1 in 100 fluorophore-labeled histone H2B proteins (unless otherwise indicated) were diluted to 1 µM nucleosome concentration in a minimal chromatin dilution buffer. For Tris-chloride conditions with glycerol this buffer was (25 mM Tris•Cl, pH 7.5, 5%[w/v] glycerol). For Tris-chloride conditions without glycerol this buffer was (25 mM Tris•Cl, pH 7.5). For Tris-acetate conditions this buffer was (25 mM Tris•Acetate, pH 7.5). For Tris-glutamate conditions this buffer was (25 mM Tris•Glutamate, pH 7.5). For PIPES-KOH conditions this buffer was (20 mM PIPES•KOH, pH 6.8). Diluted nucleosomal arrays were incubated for 5 minutes at room temperature in these minimal chromatin dilution buffers before adding 1 volume of minimal phase separation buffer. For Tris-chloride conditions the predominant minimal phase separation buffer was (25 mM Tris•Cl, pH 7.5, 300 mM NaCl, 2 mM MgCl2, 5%[w/v] glycerol). For Tris-acetate conditions the predominant minimal phase separation buffer was (25 mM Tris•Acetate, pH 7.5, 300 mM potassium acetate, 2 mM magnesium acetate). For Tris-glutamate conditions the predominant minimal phase separation buffer was (25 mM Tris•Glutamate, pH 7.5, 300 mM potassium glutamate, 2 mM magnesium glutamate). For PIPES-KOH conditions the predominant minimal phase separation buffer was (140 mM PIPES•KOH, pH 6.8, 2 mM MgCl2, 2 mM EGTA). Supplements to buffer composition were added to these phase separation buffers, with BSA at 0.2 mg/mL; DTT at 10 mM; glucose at 40 mM; Glucose Oxidase and Catalase at 20 and 3.5 µg/mL (photo-crosslinking assays) or 2 and 0.35 µg/mL (all else), respectively. For varying concentrations of salt, phase separation buffers were adjusted to achieve the intended final concentration of salt indicated in figures, figure legends, or text. After adding phase separation buffer reactions were gently and thoroughly mixed and added to a PEGylated and BSA passivated (unless otherwise indicated) microscopy well using a cut pipet tip.

Unless noted differently, condensates throughout were formed in a final buffer composition of 25 mM Tris•Acetate, pH 7.5, 150 mM potassium acetate, 1 mM magnesium acetate.

##### Partial Droplet FRAP

Nucleosomal arrays with 1 in 100 fluorophore-labeled histone H2B proteins were diluted to 2 µM nucleosome concentration into a minimal chromatin dilution buffer (see compositions outlined above). Diluted nucleosomal arrays were incubated for 5 minutes at room temperature before adding 1 volume of minimal phase separation buffer (see compositions outlined above). Supplements to phase separation buffer compositions outlined above, as indicated in text or Figure Legends, were BSA at 0.2 mg/mL; DTT at 10 mM; glucose at 40 mM; Glucose Oxidase and Catalase at 2 and 0.35 µg/mL, respectively. After adding phase separation buffer, reactions were gently and thoroughly mixed and added to an mPEGylated but not BSA passivated microscopy well using a cut pipet tip. 1 to 2 hours after condensate formation, the central third or less of the condensate by area was photobleached to ∼50% pre-bleach fluorescence intensity using 488 nm light on a laser scanning confocal microscope. Fluorescence recovery after photobleaching was measured at regular intervals using time-lapse fluorescence confocal fluorescence microscopy.

Unless noted differently, condensates in FRAP assays were formed in a final buffer composition of 25 mM Tris•Acetate, pH 7.5, 150 mM potassium acetate, 1 mM magnesium acetate.

##### Whole Droplet FRAP

Nucleosomal arrays with 1 in 100 fluorophore-labeled histone H2B proteins were diluted to 200 nM nucleosome concentration into a Minimal Acetate Dilution Buffer (25 mM Tris•Acetate, pH 7.5). Diluted nucleosomal arrays were incubated for 5 minutes at room temperature before adding 1 volume of Whole Droplet FRAP Buffer (25 mM Tris•Acetate, pH 7.5, 300 mM potassium acetate, 2 mM magnesium acetate, 0.2 mg/mL BSA, 10 mM DTT). After adding FRAP buffer, reactions were gently and thoroughly mixed and added to an mPEGylated but not BSA passivated microscopy well using a cut pipet tip. 1 to 2 hours after condensate formation, the entirety of condensates were photobleached using 488 nm light on a laser scanning confocal microscope. Fluorescence recovery after photobleaching was measured at regular intervals using time-lapse fluorescence confocal fluorescence microscopy.

##### Spinning disc confocal microscopy

Photo-crosslinking and confocal fluorescence microscopy imaging were captured on a Nikon Eclipse Ti microscope base equipped with a Yokogawa CSU-X1 spinning disk confocal scanner unit, 100 × 1.49 NA objective, and Andor EM-CCD camera.

##### Laser scanning confocal microscopy

Beside photo-crosslinking assays, all confocal fluorescence imaging was performed using a Leica SP8 confocal fluorescence microscope equipped with a resonant scanning stage, 20x dry objective, EM-CCD camera, and FRAP module. FRAP was performed using a non-resonant line scanning stage.

#### Imaging Condensates in the Narlikar Lab

##### Phase Separation of Nucleosomal Arrays

Nucleosomal arrays with 46 base pair internucleosome linker DNA length and no fluorophore label, assembled in the Narlikar lab, were diluted at 1 µM nucleosome concentration in a minimal chromatin dilution buffer. For Tris-chloride conditions with glycerol this buffer was (25 mM Tris•Cl, pH 7.5, 5%[w/v] glycerol). For Tris-acetate conditions this buffer was (25 mM Tris•Acetate, pH 7.5). For Tris-glutamate conditions this buffer was (25 mM Tris•Glutamate, pH 7.5). For PIPES-KOH conditions this buffer was (20 mM PIPES•KOH, pH 6.8). Nucleosomal arrays were incubated for 5 minutes at room temperature in these minimal chromatin dilution buffers before adding 1 volume of minimal phase separation buffer. For Tris-chloride conditions this buffer was (25 mM Tris•Cl, pH 7.5, 300 mM NaCl, 2 mM MgCl2, 5%[w/v] glycerol). For Tris-acetate conditions this buffer was (25 mM Tris•Acetate, pH 7.5, 300 mM potassium acetate, 2 mM magnesium acetate). For Tris-glutamate conditions this buffer was (25 mM Tris•Glutamate, pH 7.5, 300 mM potassium glutamate, 2 mM magnesium glutamate). For PIPES-KOH conditions this buffer was (140 mM PIPES•KOH, pH 6.8, 2 mM MgCl2, 2 mM EGTA). After adding phase separation buffer reactions were gently and thoroughly mixed, incubated for 30 minutes at room temperature, and added to a PEGylated and BSA passivated microscopy well using a cut pipet tip.

##### Bright-field Light Microscopy

Images were captured on a Widefield Nikon Ti inverted microscope base equipped with a Nikon DS-Qi2 monochrome camera and Plan Apo 40x/0.95 objective with 1.5x magnification booster. Time-lapse images were recorded over 3 minutes at 1 second intervals across 1608×1608 pixels with a pixel resolution of 122 nm x 122 nm.

#### Imaging Condensates in the Gerlich Lab

##### Phase Separation of Nucleosomal Arrays

Nucleosomal arrays with 25 bp internucleosome linker length and no fluorophore label, assembled in the Rosen lab, were diluted at 1 µM nucleosome concentration in Minimal Acetate Dilution Buffer (25 mM Tris•Acetate, pH 7.5) and incubated for 5 minutes at room temperature. Phase separation was induced by the addition of 1 volume of Minimal Acetate Phase Separation Buffer (25 mM Tris•Acetate, pH 7.5, 300 mM potassium acetate, 2 mM magnesium acetate) with and without supplement of 0.2 mg/mL BSA, 10 mM DTT, 40 mM glucose, 2 µg/mL Glucose Oxidase, and 0.35 µg/mL Catalase as indicated in the text. After adding phase separation buffer reactions were gently and thoroughly mixed and added to a PEGylated and BSA passivated microscopy well using a cut pipet tip.

##### Differential Interference Contrast Microscopy

After a 90-minute incubation, microscopy images were captured on an Axio Observer Z1/7 microscope equipped with a Plan-Apochromat 20x/0.8 objective and Hamamatsu Orca Flash 4 Camera at 2.2 Volts at 22-25°C. Time-lapse images were recorded over 10 minutes at 0.4 second intervals with 10 millisecond exposures across 2048×2048 pixels with a pixel resolution of 325 nm x 325 nm.

#### Condensate Photocrosslinking Assay

##### Magnesium-dependent Phase Separation of Nucleosomal Arrays

Nucleosomal arrays with a 25 bp internucleosome linker DNA length and 1 in 20 to 1 in 100 AlexaFluor 488-labeled histone H2B proteins were diluted to 1 µM nucleosome concentration in a Minimal Acetate Dilution Buffer (25 mM Tris•Acetate, pH 7.5). Diluted nucleosomal arrays were incubated for 5 minutes at room temperature in dilution buffer before adding 1 volume of Magnesium-dependent Phase Separation Buffer (25 mM Tris•Acetate, pH 7.5, 100 mM potassium acetate, 4 mM magnesium acetate) for a final buffer concentration of 25 mM Tris•Acetate, pH 7.5, 50 mM potassium acetate, and 2 mM magnesium acetate. Buffering supplements added as indicated to the Magnesium-dependent Phase Separation Buffer were BSA at 0.2 mg/mL; DTT at 10 mM; glucose at 40 mM; Glucose Oxidase at 20 µg/mL; Catalase at 3.5 µg/mL. After adding Magnesium-dependent Phase Separation Buffer, reactions were gently and thoroughly mixed and added to a PEGylated, but not BSA passivated, microscopy well using a cut pipet tip.

##### Photocrosslinking

One to two hours after addition to the well, condensates of varying were exposed to 0.37 to 2.1 milliWatts of 488 nm light for 500 or 50 milliseconds, as indicated, using a spinning disk confocal fluorescence microscope.

##### EDTA-dependent dissolution of non-crosslinked condensates

After fluorescent light exposure, 1 µL of 500 mM EDTA was added to 40 µL of solution in the microscopy well to dissipate condensates that were not crosslinked to one another.

##### Imaging photo-crosslinked condensates

One minute after the addition of EDTA, images were taken at the exact site of light exposure. If photocrosslinking occurred, as indicated by light-dependent persistence of EDTA-resistant condensates, images were taken adjacent to the crosslinked condensate scar (e.g., left, right, up, down).

### Quantification and Statistical Analyses

Statistical tests performed on experimental data and their representations are noted in figure legends. Image analysis was performed using ImageJ (Version 1.53) (Schneider et al., 2012). Unless otherwise described, equivalent brightness and contrast were used when depicting microscopy images in each panel. In general, microscopy data processed by ImageJ was graphed using the R Statistical Package(Team, 2013).

#### Determining Inverse Capillary Velocity from Condensate Fusion

##### Image Processing

In ImageJ, microscopy images were flatfield corrected using a gaussian blur and set to a threshold to find intrinsic chromatin condensates in each image. Particles were then analyzed, outputting a unique identifier, spatial information (e.g., XM, YM, Circularity, etc.), and Slice number for each condensate in each image.

##### Identifying condensate fusion events

Using a custom script within the R Statistical Package (available upon request), individual condensate tracks were generated across the time-lapse by finding the nearest condensate (≤ 10 µm^2^) between each frame. Bona fide fusion events were found among these condensate tracks by identifying two condensate tracks that merge coinciding with a sudden increase condensate area and aspect ratio that decreases exponentially over time. The fidelity of many of the identified condensate fusion events were manually verified in microscopy images.

##### Calculating inverse capillary velocity

For each condensate fusion event, the characteristic relaxation time (*τ*) was extracted from fit of aspect ratio (AR) over time (t) to *AR* = 1 + (*AR_init_* − 1) · *e^-t/τ^*, where *AR_init_* is the initial aspect ratio following the onset of fusion. The diameter of each condensate, before and after fusion, was calculated using the area output (in µm^2^) from ImageJ. Inverse capillary velocity was then calculated from the linear fit of *τ* and the combined pre-fusion condensate diameters across two biological replicates. The mean and standard deviation of these replicates is represented graphically in this manuscript.

#### Determining Condensate Diameters over time

##### Image Processing

In ImageJ, 5 or more microscopy images for each time point in each buffer were thresholded by fluorescence intensity to find chromatin condensates. Particles were then analyzed, outputting a unique identifier, spatial information (e.g., XM, YM, Circularity, etc.), and Slice number for each condensate in each image.

##### Calculating Condensate Diameters

Using the R Statistical package, the diameter for each identified condensate was calculated using the area output (in µm^2^) from ImageJ. These diameters were then plotted as a notched boxplot and the statistical differences between time points determined using the student’s t-test.

#### FRAP Quantitation

##### Image Processing

Image analysis was performed using ImageJ. Using unbleached condensates as a control, photobleaching was corrected across the time course using a ratio-metric technique.

##### Partial Droplet FRAP analysis

For each image in each condensate, a 20-pixel wide line plot was calculated across the condensate at the bleached locus. Partial droplet FRAP analysis was completed for each experiment using these line plots and a custom script in R (available upon request). For partial droplet FRAP of condensates composed of dodecameric nucleosomal arrays in different buffered salt solutions in Figure 2, fluorescence recovery within the bleached region was determined relative to post-bleach mean intensity. (Note: we had previously determined that these condensates mix internally over the course of fluorescence recovery (Gibson et al., 2019)). For partial droplet FRAP of all other condensates, fluorescence recovery within the bleached region was determined relative to the normalized max intensity signal of each line plot. This computational strategy is used to measure for internal mixing after photobleach (i.e. return to homogeneity) independent of differences in fluorescent molecule influx from solution between constructs and solution conditions.

##### Whole Droplet FRAP analysis

Whole droplet FRAP was determined from the mean corrected fluorescence intensities of condensates before and after photobleach.

#### Quantitation of Condensate Movement

##### Image Processing

In ImageJ, condensates were identified across 4 technical replicates per condition from 2-minute-long time-lapse microscopy acquisition of 500 milliseconds per frame. For each time point in each condition and each replicate images were set to a threshold by fluorescence intensity to find chromatin condensates. Particles were then analyzed, outputting a unique identifier, spatial information (e.g., XM, YM, Circularity, etc.), and Slice number for each condensates in each image.

##### Condensate Tracking

Using a custom script in R (available upon request), individual condensate tracks were generated across the time-lapse by finding the nearest condensate (≤ 10 µm^2^) between each frame. Custom-generated graphical depictions of condensate-centered and relative trajectories can be found in Figure 4.

##### Calculating mean squared displacement versus lag time

Condensate tracks were then segregated into 15 second segments, with 30 condensate positions per track per segment, for each 500-millisecond window. The mean squared displacement between the initial position of each segment and the lagged time positions thereafter was determined. Mean squared displacement of from all segments from all condensates between 4 and 8 microns in diameter were used to calculate the mean squared displacement versus lag time and diffusion coefficient for each condensate.

## Data and Code Availability

All custom code is available from the authors upon request.

